# Unlocking the potential of *Gordonia rubripertincta* in syngas fermentation for carbon monoxide bioconversion into carotenoids

**DOI:** 10.64898/2026.05.04.722808

**Authors:** Gayathri Vemparala, Gangagni Rao Anupoju, Thenkrishnan Kumaraguru

## Abstract

Fermentation of C1 gases is an emerging technology where waste gases are bio converted into value-added products. This study navigates the gas fermentation potential of *Gordonia rubripertincta* to produce carotenoids. The crucial carbon monoxide dehydrogenase (CODH) enzyme, necessary for gas uptake by the microbe, was found to be present in *G. rubripertincta* through blastp on NCBI website. The organism was then used for gas fermentation experiments in a continuous stirred tank reactor (CSTR) in different modes of reactor operation resulting in the production of about 500 mg pigment/g WCW (wet cell weight). Two important reactor parameters, molybdenum content and pH, were optimized for enhanced carotenoid production. Overall, *G. rubripertincta* was observed to be an efficient candidate organism for C1 gas fermentation.

**KEY HIGHLIGHTS:**

- *Gordonia rubripertincta* synthesises aerobic carbon monoxide dehydrogenase enzyme.
- It is a potential gas fermenting microbe that gives carotenoids as product.
- The gas uptake efficiency of the microbe is more in fed-batch discontinued mode.
- In FB-D, the resultant carotenoids are 500±9 mg/g wet cell weight (WCW).
- Mo/pH of 20 mg/7.0 resulted in highest carotenoids, i.e., 134±41 mg/g WCW.

**GRAPHICAL ABSTRACT:** 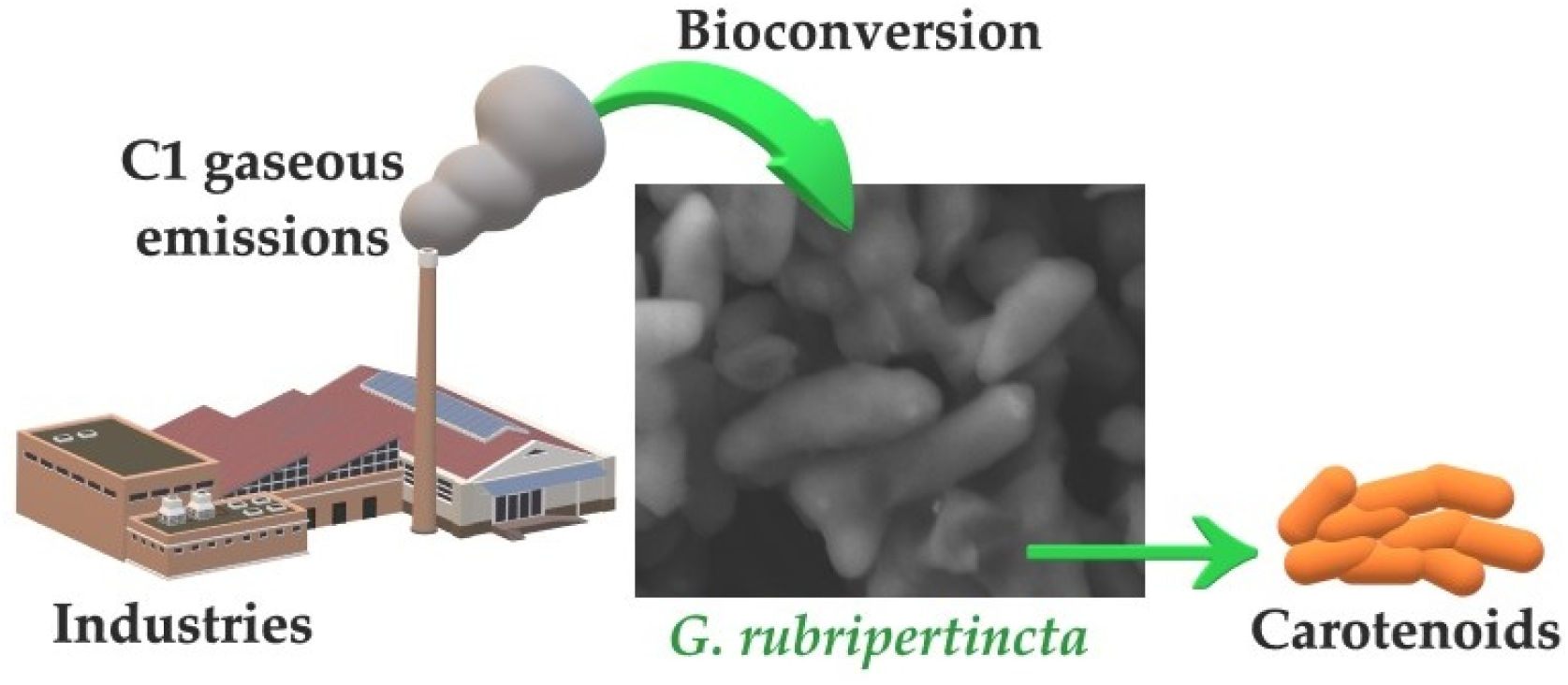

## 1. INTRODUCTION

Syngas fermentation is a microbial process where mixture of C1 gases that are potent greenhouse gases usually emitted from steel mills, chemical manufacturing industries and coal based thermal power plants are valorized into useful products.^1–5^ Bioconversion of these gases can prove to be eco-friendly as these gases are pollutants causing global warming, climate change, floods, polar ice melting, increase in sea levels, thus effecting life on earth.^6–8^

Therefore, many attempts are being made to achieve circular bioeconomy with reduced pollution and its effects. The present physicochemical methods being employed for the utilization of gaseous feedstock into value-added products are Fischer-Tropsch process (FTP)^9^, methanol-to-gasoline (MTG) process developed by Mobil Oil Corporation, Sabatier-Senderens process to synthesize hydrocarbons by hydrogenating carbon monoxide, methanol and dimethyl ether production where dimethyl ether has applications in producing, gasoline-range hydrocarbons to olefins etc.^10^ Another route of gas pollutants utilization is the biological way where microorganisms are usually employed for fermentation of C1 feedstock to produce value-added products.^11,12^ Currently, the products being synthesized via the gas fermentation route are organic acids, alcohols, polymer precursors, etc.^13–16^

The microorganisms that are currently under use include mostly bacteria and microalgae.^17,18^ Microalgae such as *Chlorella, Spirulina, Dunaliella, Cyanobacteria*, are photosynthetic microbes which can take up gaseous CO_2_ and imbibe the carbon into its metabolism.^19^ Microalgae are well known store houses of various metabolites like vitamins, carotenoids, fatty acids like docosahexanoic acid (DHA), eicosapentaenoic acid (EPA), and the whole cell can be used as single cell protein (SCP) supplement.^20^ Thus, microalgae can be one good choice for the fermentation of C1 feedstocks. Although, these pigments are produced by prokaryotes and eukaryotes, a bacterium was chosen for this study as bacteria are easy to cultivate and maintain at lab and industrial scales than eukaryotes, they finish their life cycle in few hours to very few days decreasing the industrial down time, and they are versatile in utilizing different C1 gases like CO, CO_2_, H_2,_ whereas microalgae can take up only CO_2_.

Bacteria employ a different path for taking up gaseous carbon as food source. There is a category of bacteria known as acetogens that can perform gas fermentation via Wood-Ljungdahl pathway (WLP).^21^ Some genera include *Acetobacterium, Clostridium, Butyribacterium, Moorella*.^2^ To produce acids and alcohols through gas fermentation mostly bacteria are used as catalysts. Another category of bacteria that can perform gas fermentation are carboxydotrophs. Carboxydotrophs include both aerobic and anaerobic types of bacteria.^22,23^ The aerobic carboxydotrophs synthesize aerobic carbon monoxide dehydrogenase (CODH) enzyme that is key for the uptake of C1 gases.^24,25^ *Gordonia* is one such genus having aerobic carboxydotrophic species. *Gordonia rhizosphera* was reported to be growing in areas rich in CO gases indicating its ability of assimilating C1 gaseous substrates.^26^ As there is probability that *Gordonia* species can ferment C1 gases and produce carotenoids, an existing *Gordonia rubripertincta* organism was chosen for this work.

On the other hand, many *Gordonia* species were reported to be producers of various important metabolites like vitamins, enzymes, carotenoids, surface active compounds etc.^27^ Carotenoids are secondary metabolites produced by many microorganisms, fungi, algae, plants, and animals. They are yellow to red color pigments that have prophylactic activity and are important compounds for pharmaceutical, cosmetic and food industries.^28^ Carotenoids are useful as nutraceuticals to prevent diseases or provide supplements required by humans or animals.^29^ Previously, *Gordonia jacobeae* was reported to be a canthaxanthin producer.^30^ *Gordonia alkanivorans* was shown as a potential desulfurizer and it also produces Canthaxanthin, astaxanthin carotenoids^31,32^. *Gordonia ajoucoccus, G. terrae, G. amicalis* were all studied previously and identified to produce γ-carotene and its derivatives.^27,33,34^

However, none of the previous studies focused on exploring the C1 gas fermentation capacity of *G. rubripertincta* and hence, it was chosen as the candidate organism for the case study of carotenoids production via gas fermentation route.

In this study, the CODH enzyme presence in *Gordonia rubripertincta* was confirmed by blastp on NCBI website. Once the enzyme synthesis was confirmed, the organism’s ability to take up gaseous carbon as feedstock was tested practically in a Continuous Stirred Tank Reactor (CSTR) with sole CO as carbon and energy source. The growth pattern and carotenoids quantity were analyzed after each experiment. Reactor parameters such as mode of operation, molybdenum content and pH were optimized to enhance the carotenoid production.

## 2. MATERIALS AND METHODS

### 2.1 Microbial culture collection and maintenance

The bacterium under study, *Gordonia rubripertincta*, was procured from Microbial Type Culture Collection (Accession No. 289), CSIR-Institute of Microbial Technology, Chandigardh, India. It was maintained and sub-cultured in the lab using Luria-Bertani (LB) medium, and the culture was stored under 2-4 °C. The liquid medium used for gas fermentation experiments is the Modified Basal Medium (MSM) whose composition is: Yeast extract 1 g/L, Mineral solution 25 mL/L, Trace metal solution 10 ml/L, Vitamin solution 10 mL/L, Resazurin 1 mL/L. Mineral stock solution contains, Sodium Chloride 80 g/L, Ammonium Chloride 100 g/L, Potassium Chloride 10 g/L, Potassium monophosphate 10 g/L, Magnesium sulphate 20 g/L, Calcium chloride 4 g/L. Trace metal stock solution is made up of, Nitriloacetic acid (NTA) 2 g/L, Manganese sulphate 1 g/L, Ferrous Ammonium Sulphate 0.8 g/L, Cobalt chloride 0.2 g/L, Zinc sulphate 0.2 g/L, 20 mg each of Copper (II) chloride, Nickel chloride, Sodium molybdate, Sodium selenate and Sodium tungstate.^13^

### 2.2 Computational identification of CODH enzyme synthesis by *G. rubripertincta* using blastp

For this analysis, protein(s) information existing on the National Centre for Biotechnology Information (NCBI) website was used. The CODH protein sequence of *Gordonia rhizosphere* was given as the query sequence (Query ID: WP_006329678.1) and was compared with the whole proteins of *Gordonia rubripertincta*. A blastp was run to identify if the organism being studied, *G. rubripertincta*, contains the similar protein sequence based on sequence alignment and a similarity depicting phylogenetic tree was also constructed.

### 2.3 Initial batch gas fermentation experiment with *G. rubripertincta*

In all the gas fermentation experiments, sole and pure carbon monoxide (CO) was supplied as the carbon and energy source to the microbe. The fermentor used was Continuous Stirred Tank Reactor (CSTR) with MSM **(composition given in Section 2.1)** supplying trace elements, essential vitamins, and minerals to the microbe. The working volume of the reactor was 2 L which was added with MSM and sterilized. Later, 10% v/v *G. rubripertincta* culture was added. The reactor was set for fermentation with 1 L headspace filled with CO gas. The RPM maintained was 500 at 25 °C. The fermentation was carried out for 5 d, a period set based on the growth pattern of the organism, and everyday liquid samples were collected to analyze the growth pattern in terms of OD at 600 nm using a UV-VIS spectrophotometer and final carotenoids quantification.

### 2.4 Fed-batch (FB) Fed-batch discontinued (FB-D) gas fermentation

The reactor conditions were exactly same as in batch fermentation, 2 L MSM with 10% v/v microbial culture, 500 RPM, 25 °C temperature, except for gas sparging. The CO gas was supplied 1 L every day for the whole fermentation period of 5 d bringing the total gas sparged to 6 L with regular collection of liquid samples. The samples were analyzed for growth and carotenoid production. Later, a modification was made in the fed-batch technique by discontinuing gas sparging after 2 d so that the total gas sparged was 3 L. CO gas supply was given only initially, after 24 and 48 h in the discontinued fed-batch fermentation. This was also carried out for 5 d and liquid samples were regularly collected for analysis.

### 2.5 Molybdenum and pH optimization

Experiments were designed in a 2^2 factorial design fashion with two variations in each of the factors, Mo and pH, and the results were statistically analyzed using R software. As the MSM already contains molybdenum in the form of 20 mg sodium molybdate, an additional 10 and 20 mg were given to increase the quantity of Mo to 30 mg and 40 mg, and check its effect on growth and pigmentation. Similarly, pH of 6.5 and 7.0 were tested. The optimization was done for both the factors in combination, so, four experiments were carried out with 30 mg/6.5 pH (R1), 30 mg/7.0 pH (R2), 40 mg/6.5 pH (R3), and 40 mg/7.0 pH (R4). The fermentation was carried out for 5 d at 500 RPM, 25 °C temperature and 1 L CO gas was sparged only on the initial day with regular sampling for growth and carotenoids analysis.

### 2.6 Solvent extraction for quantitative estimation of carotenoids

The carotenoids *G. rubripertincta* produces are completely intracellular and therefore, solvent extraction method was employed to bring them out of the cell and quantify. Methanol solvent was used for all the extractions. To 1 g of wet cell weight (WCW) 2 mL of methanol was added and vortexed for a minute. Then it was centrifuged at 6000 RPM for 5 min and the upper pigment containing solvent layer was collected. The extraction process was repeated for 4 times and all the carotenoids dissolved in methanol were pooled together. To quantify, the carotenoids solution was taken in a pre-weighed clean and dry round bottom flask, solvent was dried using a vacuum rotavapor, and the carotenoids remaining at the end were weighed. The same carotenoids extraction and quantification procedure was followed for all the reactor experiments including controls and duplicates.

### 2.7 Statistical analysis

All the experiments were carried out in duplicates. The obtained results were statistically analyzed to verify their accuracy and reliability which was performed using MS Excel and R software. One-way ANOVA for batch, fed-batch and fed-batch (discontinued) were done using MS Excel while Two-way ANOVA for Mo and pH combination experiments was done on R along with building a regression model, contour, and residual plots.

## 3. RESULTS AND DISCUSSION

### 3.1 Computational identification of CODH enzyme synthesis by *G. rubripertincta* using blastp

Previously, CODH was reported to be an enzyme essential for CO oxidation by carboxydotrophs. In *Pseudomonas carboxydovorans* CODH was reported to be present in either cytoplasm or cell membrane.^24^ The CODH in *Oligotropha carboxidovorans* is an enzyme with one large and two small subunits and the larger subunit has a Mo/Cu active site.^25^ *Gordonia rhizosphera* was found to be a CO oxidizer that contains CODH enzyme and the large subunit (CoxL) is of form I.^26^ It can be understood that CODH enzyme is very important for an organism to uptake CO gas and assimilate it. Thus bioinformatics studies were conducted to find out the presence of CODH in the microbe under study, *Gordonia rubripertincta*.

The blastp run revealed the presence of CODH enzyme in *G. rubripertincta* similar to *G. rhizosphera*. The molybdopterin-dependent oxidoreductase (Accession: WP_005196015.1) present in *G. rubripertincta* showed the highest query cover of 99%. It is a 927 amino acid protein containing both small (CoxS) and large subunit (CoxL) of the enzyme **(Supplementary information: blastp results (DOC))**. The phylogenetic tree depicts the enzymes from both organisms are very closely related and might have a similar function **(Fig.1)**. The bacterium, *Gordonia rubripertincta*, synthesizes CODH enzyme which is very similar to the one produced by *Gordonia rhizosphera*, which theoretically proves that the organism under study can perform C1 gas fermentation.

**Fig. 1.**
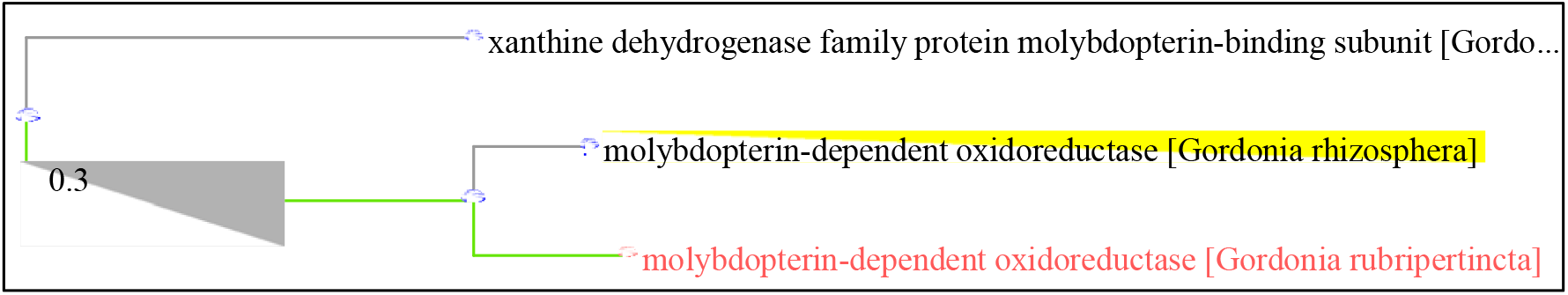
Phylogenetic tree showing the closeness between *G. rhizosphera* and *G. rubripertincta* CODH enzymes.

### 3.2 Initial batch gas fermentation with *G. rubripertincta*

The batch gas fermentation was carried out with two controls to confirm the uptake of CO by *G. rubripertincta*. One is the MSM without CO sparging, i.e., devoid of carbon and energy source, and second is the LB broth, a synthetic medium with all required nutrients including carbon dissolved in the medium. The growth pattern was monitored in all the three reactors during the complete 5 d of fermentation which can be seen in **Fig.2**. Among the three shows that LB broth has resulted in highest biomass accumulation followed by MSM+CO and MSM. LB is a medium with all required macro and micronutrients dissolved in water which will be readily available for the microbe to absorb for its maintenance, survival, and growth. Due to the highly favorable conditions, growth and carotenoid production is higher in the LB broth than MSM and MSM+CO. The organism also goes through log, stationary and decline phases when grown in LB medium. Between MSM and MSM+CO, lower biomass concentration is in MSM without CO. The difference in growth is very minimum since the start towards the end of fermentation. In the MSM, the liquid in which the microbe is added contains only trace elements, minerals and vitamins required for the growth of *G. rubripertincta*. The crucial element, carbon, is present in very minute quantity in the form of yeast extract. The carbon that is necessary for the microbe must be taken up from CO gas that is supplied. As there is no CO in MSM reactor, the change in growth is quite less. In MSM+CO the growth is higher compared to MSM showing the positive effect of CO on biomass increase. In MSM and MSM+CO also different phases of growth can be seen. In MSM+CO reactor, 1L CO was sparged at the start of fermentation and it is clear from the results that biomass concentration is higher in comparison to no CO reactor. This proves, the mass transfer of CO to liquid was possible in a CSTR and that the organism is potential enough to use a gaseous carbon source. The organism produced highest amount of carotenoids when grown in LB, followed by MSM+CO and MSM, like the growth pattern. The amounts of carotenoids produced in LB, MSM+CO, MSM are 140±20 mg, 114±17 mg, 92±4 mg/g WCW respectively. The biomass concentration or the carotenoids production is lesser in MSM+CO than in LB. This can be attributed to the fact that, the CO sparged might not be enough for the initial biomass to grow, or not every molecule of CO sparged might have reached the organism due to less solubility of CO in water, or some CO might escape from the reactor while or after sparging from the 0.2 μm membrane filter closed air outlet of the reactor. Nevertheless, the fact that CODH is produced by *G. rubripertincta* and is efficient in performing C1 gas fermentation was established. This proves that *G. rubripertincta* can utilize gaseous CO as substrate and release carotenoids as value-added product. The results were found to be statistically significant with a p-value of <0.05.

**Fig. 2.**
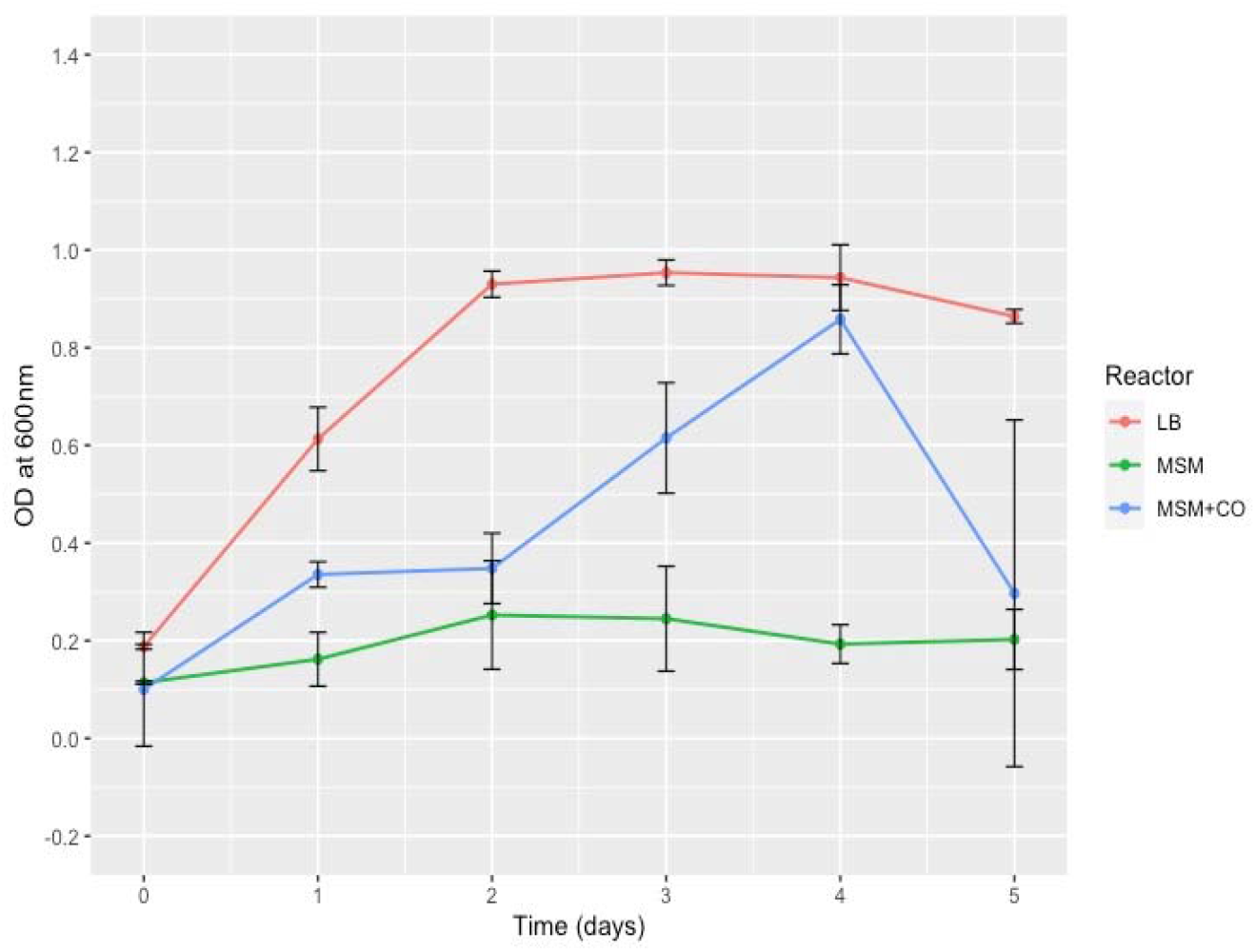
Growth curve of *G. rubripertincta* in batch gas fermentation experiments.

In continuation to the current study, the pigments produced by *G. rubripertincta* were extracted and the carotenoids were analyzed through High Resolution Mass Spectrometry (HRMS) (unpublished work).

### 3.3 Fed-batch (FB) Fed-batch discontinued (FB-D) gas fermentation

To maximize the carotenoid yield, fed-batch fermentation was tried expecting more gaseous uptake, more biomass, and carotenoids. Here, a batch operated reactor was used as control. Fed-batch mode of reactor operation was employed, where 1L of CO gas was fed everyday expecting to bring down the substrate deficiency, if there was any. From **Fig.3A**, it can be noticed that, biomass concentration has increased substantially in comparison with batch mode reactor and the total carotenoids concentration has almost doubled up, shown in **Fig.3B**, demonstrating the efficiency of gas fermentation of the microbe under study can be increased when used in a CSTR with FB mode of operation. In FB, *G. rubripertincta* continued to be in exponential phase even on the last day of fermentation. This might be due to the regular gas feed that let high biomass concentration to form. The carotenoids are secondary metabolites, synthesized by the bacterium in the stationary phase of growth. Bringing the bacterium to stationary phase was assumed to increase the carotenoid production. This must be achieved with no compromise on the high biomass accumulation as more biomass might give more amount of product. Therefore, the fed-batch mode was restricted to 2 d, i.e., gas supply was cut off after 0 d, 1 d, and 2 d of fermentation, termed here as fed-batch (discontinued) mode of reactor operation. The growth pattern as can be seen from **Fig.3A**, in FB-D reactor the biomass concentration remained like FB and sometimes even higher a little, with a stationary and decline phase in the growth. According to **Fig.3A** in the batch and FB-D operated reactors, the microbe has gone through log, stationary and decline phases. Therefore, it can be concluded that different modes operation result in different phases of growth of the microorganism. Nevertheless, total carotenoids produced in terms of concentration, is highest in FB-D followed by FB and batch modes which is given in **Fig.3B**. The total amount of carotenoids produced in FB-D, FB, and batch modes are 500±9 mg, 447±11 mg, and 217±11 mg/g WCW respectively. It can thus be established that FB and FB-D modes result in higher quantities of carotenoids than batch mode. Also, more gas can be utilized in one cycle of fermentation when FB or FB-D are employed. It can be concluded that gas feed at regular intervals, increases the uptake of CO by the organism which showed up in terms of higher biomass and carotenoid production. The results were found to be statistically significant with a p-value of <0.05.

**Fig. 3A.**
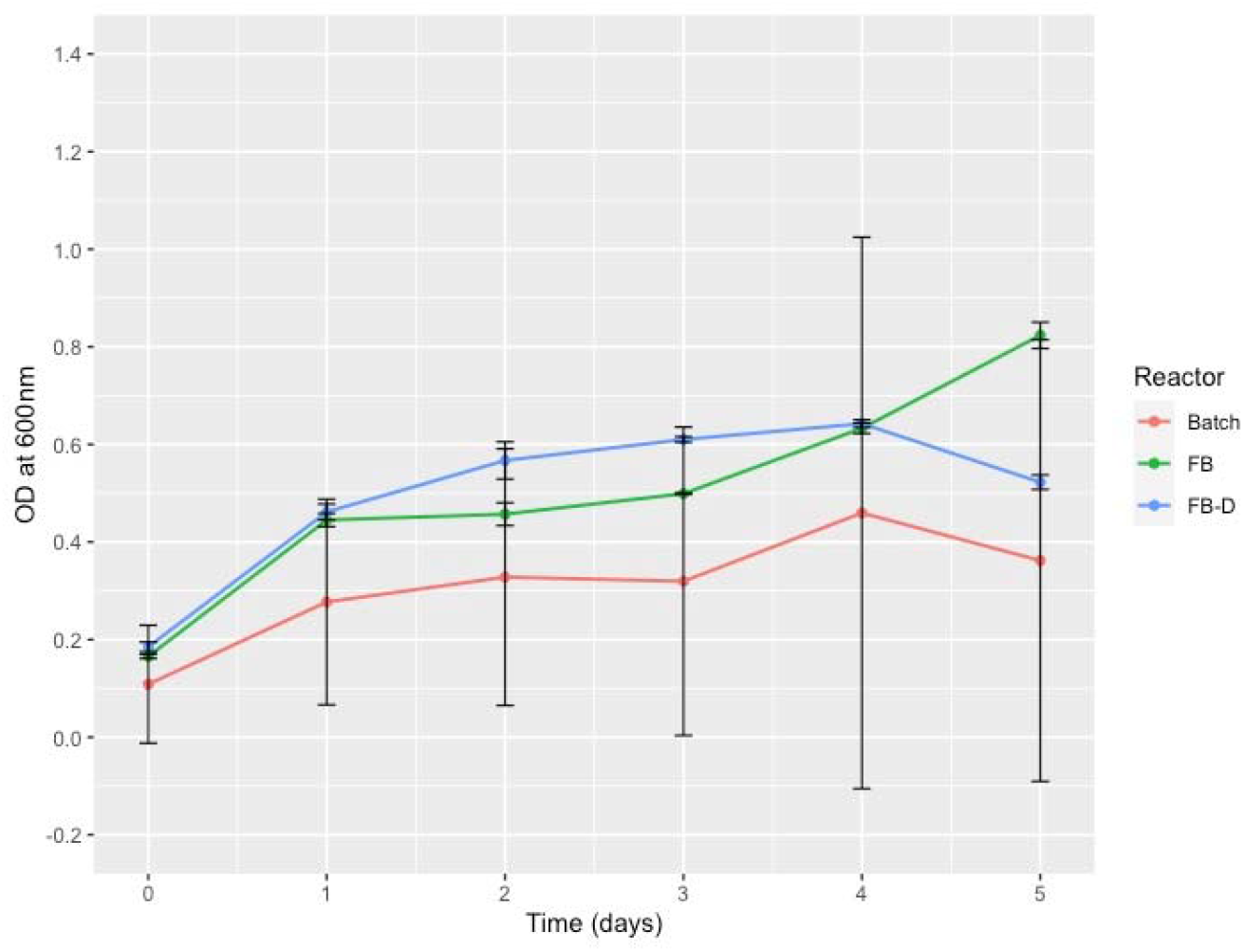
Growth curve of *G. rubripertincta* in FB and FB-D mode of gas fermentation.

**Fig. 3B.**
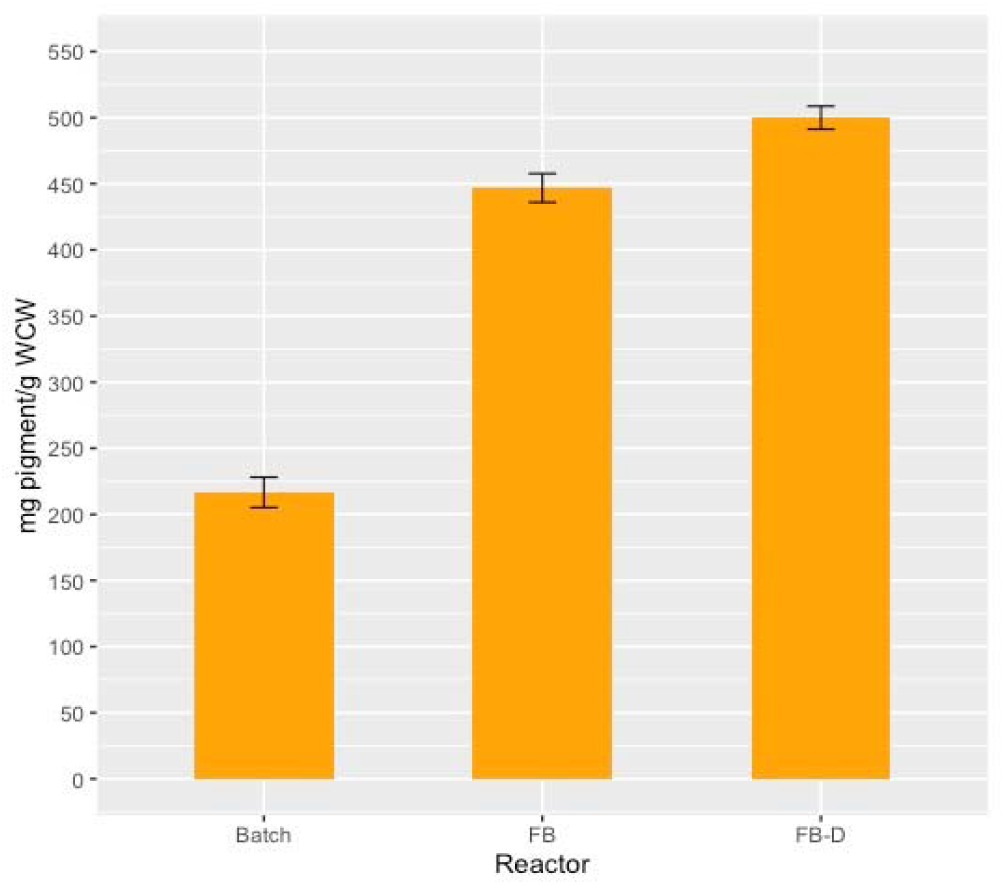
Total carotenoids produced by *G. rubripertincta* in FB and FB-D mode of gas fermentation.

### 3.4 Molybdenum and pH optimizations

The key enzyme CODH is a molybdopterin-dependent oxidoreductase, for which molybdenum is an important cofactor for its effective action in CO uptake by the microbial cell.^35^ Also, pH varies the solubility of gases in liquids increasing or decreasing its availability for the microorganism to utilize it.^36,37^ Molybdenum concentration and pH were the key physicochemical parameters chosen to try and enhance the carotenoid production by assessing their combinational effect.

Among the four experiments, the growth of the bacterium is highest in R4 and lowest in R1 which can be observed from **Fig.4A**. It can be seen that the growth pattern is very much similar in all combinations. There was lag, log, stationary phases in all reactors and a prominent decline phase are present in R1 and R2. The highest biomass concentration was accumulated when Mo/pH was at the highest point, i.e., R4, showing the positive effect of increasing the parameters on biomass concentration increase. When pH was increased to 7.0 with Mo concentration kept low at 30 mg, the shift in growth is much higher compared to R1 and R3. pH might be having a bigger effect on growth than Mo. However, when both conditions were increased together, it resulted in highest concentration of biomass.

**Fig. 4A.**
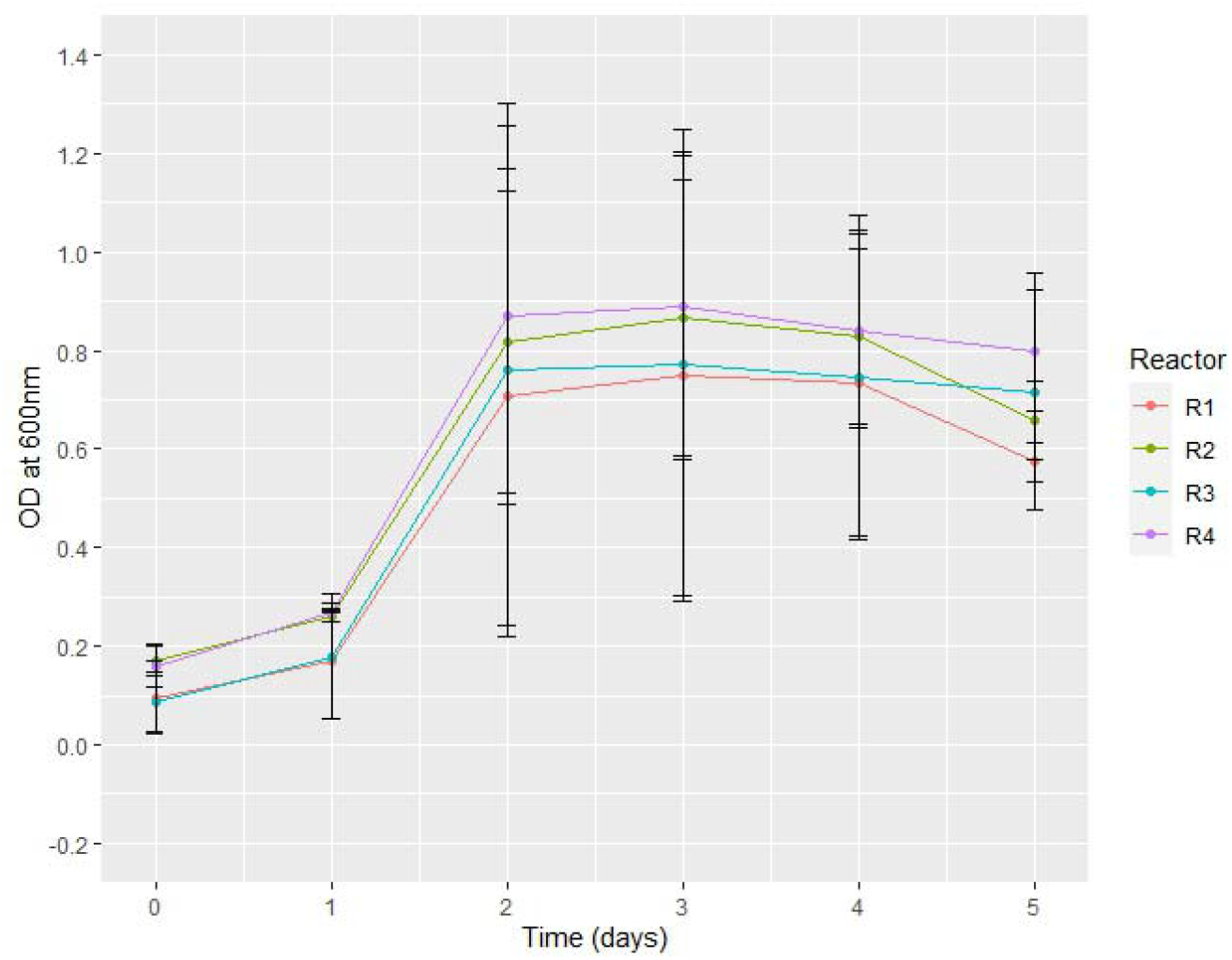
Growth curve of *G. rubripertincta* in different combinations of Mo and pH.

The total carotenoids obtained from R1, R2, R3, and R4 are 62±12 mg, 88±25 mg, 108±77 mg, and 134±41 mg pigment/ g WCW respectively and shown in **Fg.4B**. The highest carotenoids amount was obtained from the reactor R4, similar to growth curve. Contrary to the growth pattern, R3 resulted in higher pigment production than R2 pH. The carotenoids were however highest in the reactor operated at R4 and lowest in R1 same as the growth pattern. The carotenoids produced when statistically analyzed with two-factor ANOVA using R and the p-value for each factor results were <0.05.

**Fig. 4B.**
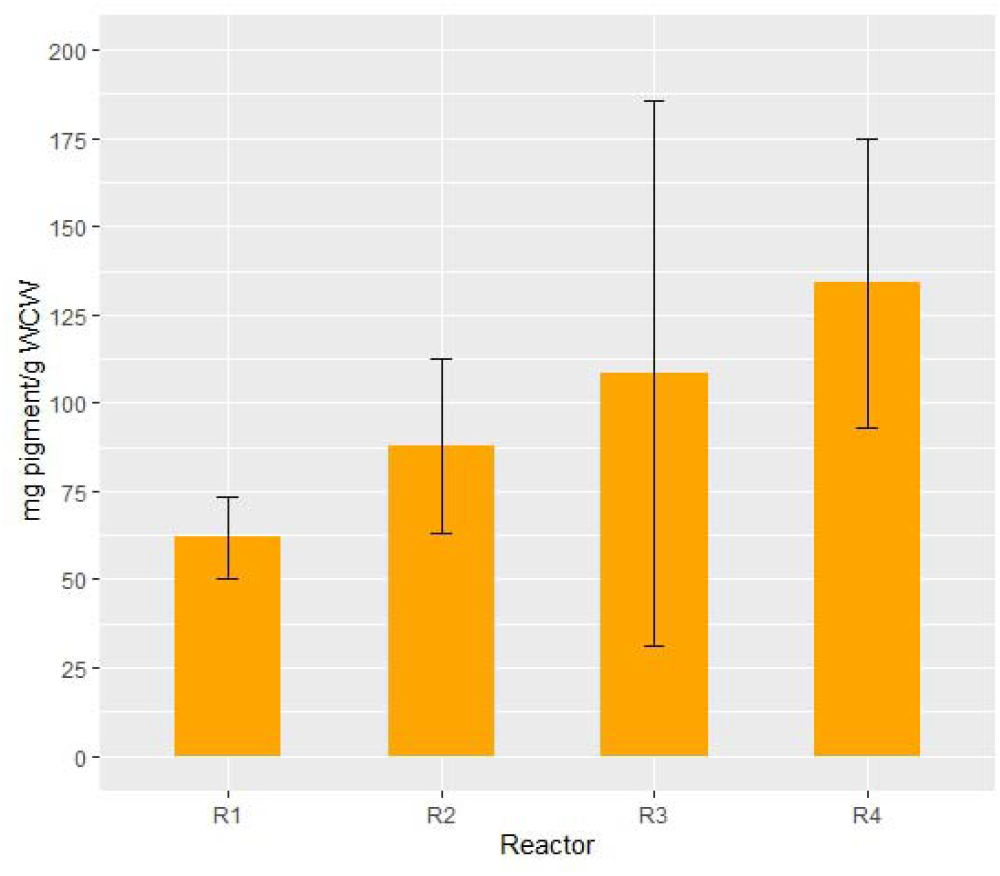
Total carotenoids produced by *G. rubripertincta* in different combinations of Mo and pH.

All the 2^2 factorial analysis done using R were tabulated and presented in **Supplementary information: DoE (XLSX)**. A regression model was built with both natural and coded variables to fit the data and obtain contour lines plot to illustrate the behavior of the yield with different Mo concentrations and pH. The R^2^ value for the regression model built based on natural variables is 89.49% indicating the precision with which the model can predict the experimental results is about 90%. The coefficient values of the regression model built on coded variables has a positive sign indicating the positive correlation between both the parameters. The coefficient for Mo concentration is 18.5, meaning, there will be about a 37% increase in the carotenoid amount when Mo is shifted from lower (30 mg) to higher (40 mg) level. Similarly, the coefficient of pH coded variable is 36, indicating there will be an approximate 72% increase in carotenoids production when pH is shifted from 6.5 to 7.0. The contour plot **(Fig.4C)** also shows increase in carotenoid production when Mo and pH were at highest levels out of the two levels tested. To check the model adequacy, residuals plot was also constructed. The residuals plot is given in **Fig.4D** showing the residuals of 8 observations of four experiments in duplicates. The residuals are randomly distributed showing the experimental procedure is good. Therefore, it is statistically also confirmed that Mo and pH variation will affect the final carotenoid production by *G. rubripertincta* using CO as feedstock.

It can thus be concluded that, *Gordonia rubripertincta* possesses the key aerobic carbon monoxide dehydrogenase (CODH) enzyme essential for assimilating C1 gases, confirmed via NCBI blast analysis. When the microorganism was used for CO gas fermentation in a CSTR, the highest pigment yield was obtained using the fed-batch (discontinued) mode of operation. The impact of Molybdenum (Mo) content, a crucial CODH cofactor, and pH, which affect gas solubility, were examined to improve carotenoid production. In summary, *G. rubripertincta* emerges as a promising candidate for converting C1 gases to carotenoids, with ongoing optimization focusing on parameters such as dissolved oxygen, temperature, and light conditions.

**Fig. 4C.**
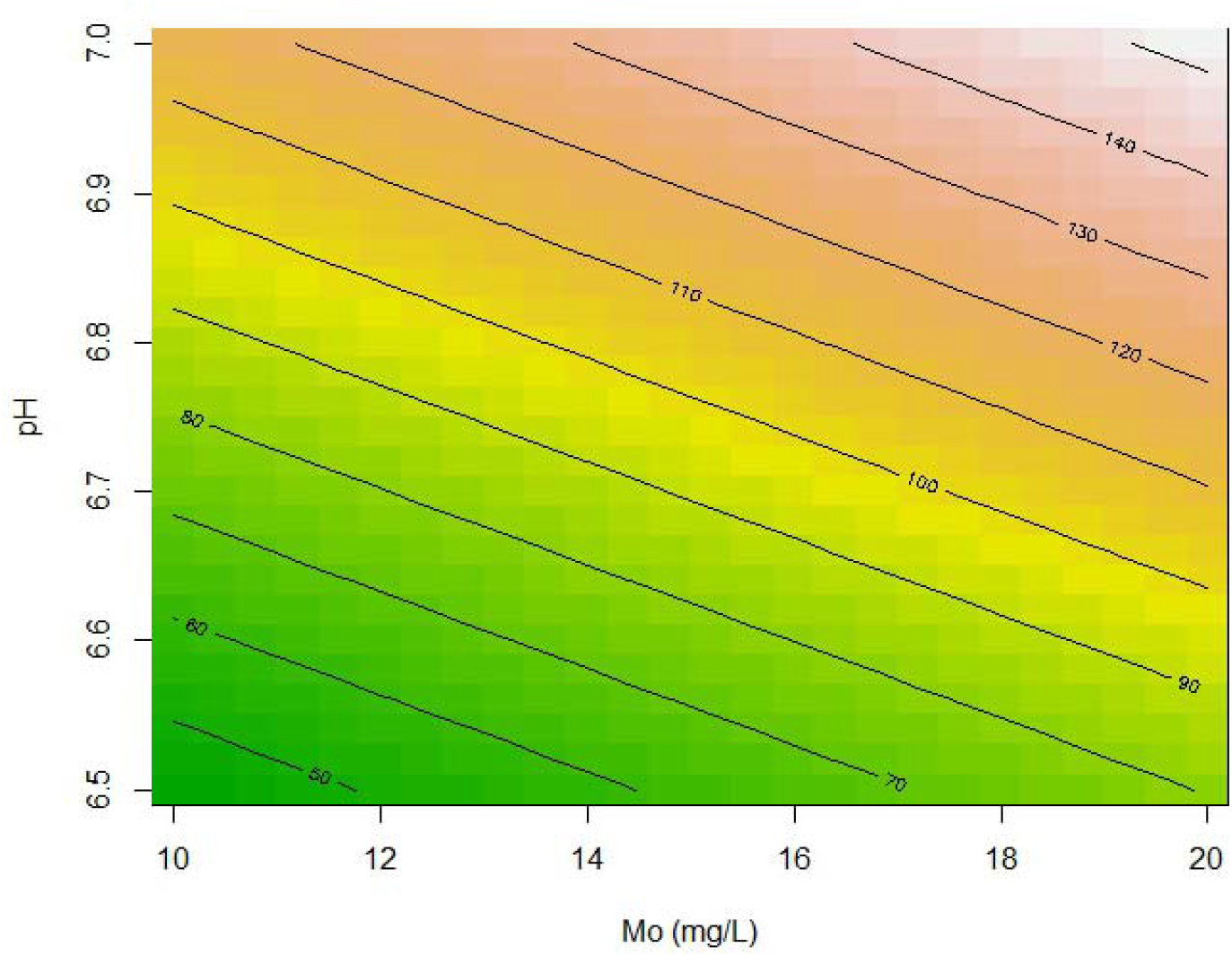
Contour plot showing the variation in carotenoid quantity with shift in Mo and pH.

**Fig. 4D.**
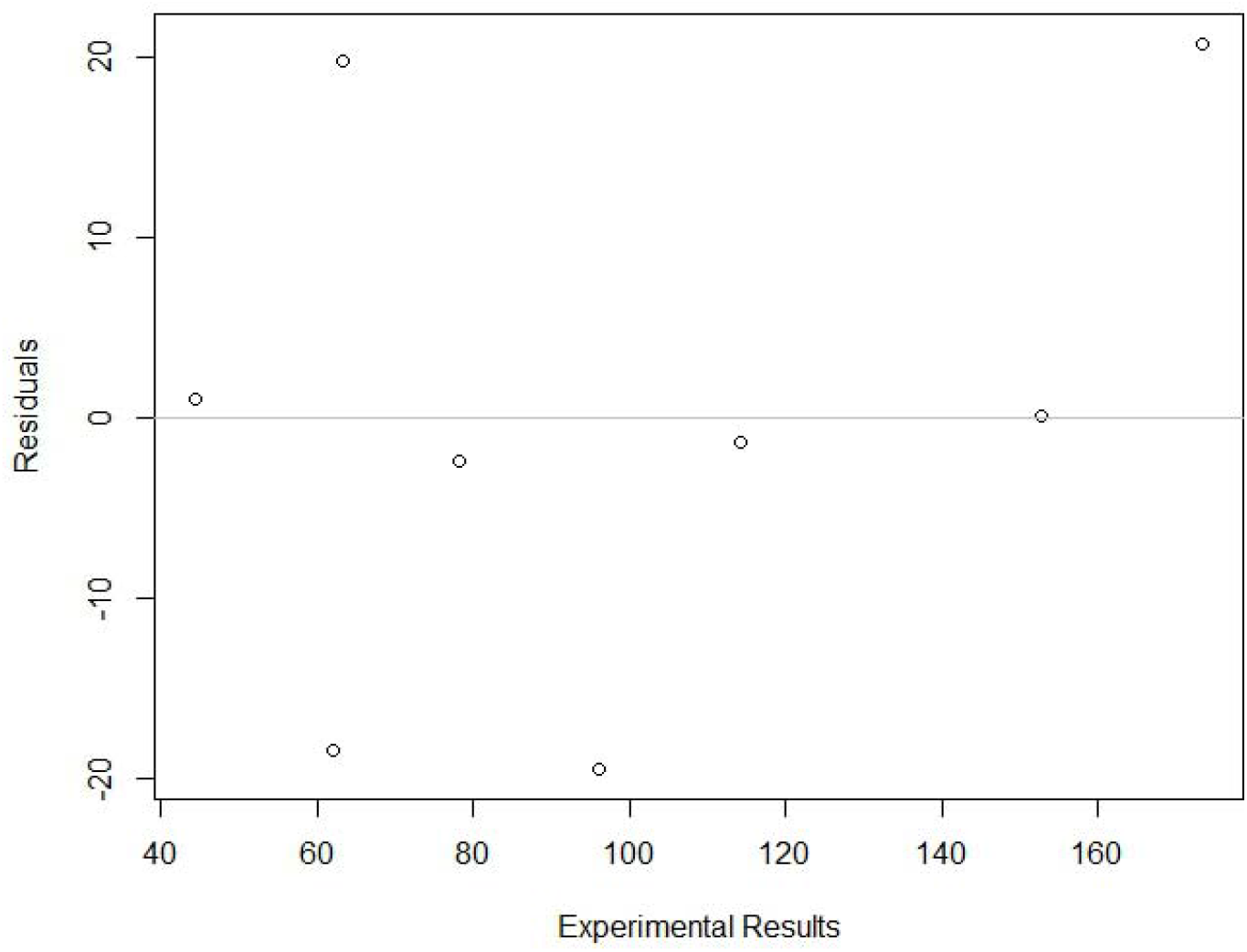
Residuals plot.

## Supporting information

blastp

DoE

## ACKNOWLEDGEMENT

The authors are thankful to The Director, CSIR-Indian Institute of Chemical Technology (IICT) for providing the facilities required for carrying out this research work (**IICT/Pubs./2024/095**). The authors are grateful to the Academy of Scientific and Innovative Research (AcSIR) for their immense support during this work. Also, sincere thanks to the Indian Council of Medical Research (ICMR) for providing Ms. Gayathri Vemparala with research fellowship (JRF-2019, HRD-22).

## REFERENCES

(1) Redl, S.; Diender, M.; Jensen, T. Ø.; Sousa, D. Z.; Nielsen, A. T. Exploiting the Potential of Gas Fermentation. Ind. Crops Prod. 2017. 10.1016/j.indcrop.2016.11.015.

(2) Liew, F. M.; Martin, M. E.; Tappel, R. C.; Heijstra, B. D.; Mihalcea, C.; Köpke, M. Gas Fermentation-A Flexible Platform for Commercial Scale Production of Low-Carbon-Fuels and Chemicals from Waste and Renewable Feedstocks. Front. Microbiol. 2016, 7 (MAY). 10.3389/fmicb.2016.00694.

(3) Monir, M. U.; Yousuf, A.; Aziz, A. A. Syngas Fermentation to Bioethanol; INC, 2020. 10.1016/b978-0-12-815936-1.00006-x.

(4) Frazão, C. J. R.; Walther, T. Syngas and Methanol-Based Biorefinery Concepts. Chemie-Ingenieur-Technik 2020, 92 (11), 1680–1699. 10.1002/cite.202000108.

(5) Fernández, Á.; Haris, N.; Abubackar, N.; Veiga, M. C. Production of Chemicals from C1 Gases (CO , - CO 2) by Clostridium Carboxidivorans. World J. Microbiol. Biotechnol. 2017, 0 (0), 0. 10.1007/s11274-016-2188-z.

(6) The greenhouse effect. bgs.ac.uk/discovering geology/climate-change.

(7) Denchak, M. Greenhouse Effect 101. NRDC stories 2019, 20.

(8) Timperley, J. The carbon brief profile: India. carbonbrief.org.

(9) Molitor, B.; Richter, H.; Martin, M. E.; Jensen, R. O.; Juminaga, A.; Mihalcea, C.; Angenent, L. T. Carbon Recovery by Fermentation of CO-Rich off Gases - Turning Steel Mills into Biorefineries. Bioresource Technology. Elsevier Ltd September 1, 2016, pp 386–396. 10.1016/j.biortech.2016.03.094.

(10) Speight, J. G. Synthetic Liquid Fuel Production from Gasification; © 2015 Woodhead Publishing Limited. All rights reserved., 2015. 10.1016/B978-0-85709-802-3.00007-2.

(11) Devarapalli, M.; Lewis, R. S.; Atiyeh, H. K. Continuous Ethanol Production from Synthesis Gas by Clostridium Ragsdalei in a Trickle-Bed Reactor. Fermentation 2017, 3 (2). 10.3390/fermentation3020023.

(12) Mohammadi, M.; Najafpour, G. D.; Younesi, H.; Lahijani, P.; Uzir, M. H.; Mohamed, A. R. Bioconversion of Synthesis Gas to Second Generation Biofuels: A Review. Renewable and Sustainable Energy Reviews. 2011. 10.1016/j.rser.2011.07.124.

(13) Fernández-naveira, Á.; Abubackar, H. N.; Veiga, M. C. Efficient Butanol-Ethanol (B-E) Production from Carbon Monoxide Fermentation by Clostridium Carboxidivorans. 2016. 10.1007/s00253-015-7238-1.

(14) Abubackar, H. N.; Veiga, M. C.; Kennes, C. Carbon Monoxide Fermentation to Ethanol by Clostridium Autoethanogenum in a Bioreactor with No Accumulation of Acetic Acid. Bioresour. Technol. 2015, 186, 122–127. 10.1016/j.biortech.2015.02.113.

(15) Fernández-naveira, Á.; Abubackar, H. N.; Veiga, M. C. Carbon Monoxide Bioconversion to Butanol-Ethanol by Clostridium Carboxidivorans□: Kinetics and Toxicity of Alcohols. 2016, 4231–4240. 10.1007/s00253-016-7389-8.

(16) Stoll, K. I.; Herbig, S.; Sauer, J.; Neumann, A.; Oswald, F. Fermentation of H 2 and CO 2 with Clostridium Ljungdahlii at Elevated Process Pressure – First Experimental Results. Chem. Eng. Trans. 2018, 64 (2), 2–7.

(17) Beigbeder, J. B.; Sanglier, M.; De Medeiros Dantas, J. M.; Lavoie, J. M. CO2 Capture and Inorganic Carbon Assimilation of Gaseous Fermentation Effluents Using Parachlorella Kessleri Microalgae. J. CO2 Util. 2021, 50 (March), 101581. 10.1016/j.jcou.2021.101581.

(18) Kassim, M. A.; Meng, T. K. Carbon Dioxide (CO2) Biofixation by Microalgae and Its Potential for Biorefinery and Biofuel Production. Sci. Total Environ. 2017, 584–585, 1121–1129. 10.1016/j.scitotenv.2017.01.172.

(19) Klinthong, W.; Yang, Y. H.; Huang, C. H.; Tan, C. S. A Review: Microalgae and Their Applications in CO2 Capture and Renewable Energy. Aerosol Air Qual. Res. 2015, 15 (2), 712–742. 10.4209/aaqr.2014.11.0299.

(20) Nethravathy, M. U.; Mehar, J. G.; Mudliar, S. N.; Shekh, A. Y. Recent Advances in Microalgal Bioactives for Food, Feed, and Healthcare Products: Commercial Potential, Market Space, and Sustainability. Compr. Rev. Food Sci. Food Saf. 2019, 18 (6), 1882–1897. 10.1111/1541-4337.12500.

(21) Bengelsdorf, F. R.; Beck, M. H.; Erz, C.; Hoffmeister, S.; Karl, M. M.; Riegler, P.; Wirth, S.; Poehlein, A.; Weuster-Botz, D.; Dürre, P. Bacterial Anaerobic Synthesis Gas (Syngas) and CO2 + H2 Fermentation. In Advances in Applied Microbiology; Academic Press Inc., 2018; Vol. 103, pp 143–221. 10.1016/bs.aambs.2018.01.002.

(22) Henstra, A. M.; Sipma, J.; Rinzema, A.; Stams, A. J. Microbiology of Synthesis Gas Fermentation for Biofuel Production. Current Opinion in Biotechnology. 2007. 10.1016/j.copbio.2007.03.008.

(23) Cantera, S.; Tamarit, D.; Strong, P. J.; Sánchez-Andrea, I.; Ettema, T. J. G.; Sousa, D.Z. Prospective CO2 and CO Bioconversion into Ectoines Using Novel Microbial Platforms. Rev. Environ. Sci. Biotechnol. 2022, 21 (3), 571–581. 10.1007/s11157-022-09627-y.

(24) Meyer, O.; Frunzke, K.; Gadkari, D.; Jacobitz, S.; Hugendieck, I.; Kraut, M. Utilization of Carbon Monoxide by Aerobes: Recent Advances. FEMS Microbiol. Rev. 1990, 7, 253–260.

(25) Gourlay, C.; Nielsen, D. J.; Evans, D. J.; White, J. M.; Young, C. G. Models for Aerobic Carbon Monoxide Dehydrogenase: Synthesis, Characterization and Reactivity of Paramagnetic MoVO(μ-S)CuI Complexes. Chem. Sci. 2018, 9 (4), 876–888. 10.1039/c7sc04239f.

(26) Hernández, M.; Vera-Gargallo, B.; Calabi-Floody, M.; King, G. M.; Conrad, R.; Tebbe, C. C. Reconstructing Genomes of Carbon Monoxide Oxidisers in Volcanic Deposits Including Members of the Class Ktedonobacteria. Microorganisms 2020, 8 (12), 1–17. 10.3390/microorganisms8121880.

(27) Sowani, H.; Kulkarni, M.; Zinjarde, S. An Insight into the Ecology, Diversity and Adaptations of Gordonia Species. Crit. Rev. Microbiol. 2018, 44 (4), 393–413. 10.1080/1040841X.2017.1418286.

(28) López, G.; Álvarez-rivera, G.; Carazzone, C.; Ibáñez, E.; Leidy, C.; Cifuentes, A. Carotenoids in Bacteria□: Biosynthesis , Extraction , Characterization and Applications. 2021, No. August. 10.20944/preprints202108.0383.v1.

(29) Souyoul, S. A.; Saussy, K. P.; Lupo, M. P. Nutraceuticals: A Review. Dermatol. Ther. (Heidelb). 2018, 8 (1), 5–16. 10.1007/s13555-018-0221-x.

(30) Villa, T. G.; Sierio, C.; de Miguel, T.; Poza, M. Isolation and Taxonomic Study of a New Canthaxanthin-Containing Bacterium, Gordon/a Jacobaea MV-1 Sp. Nov. Int. Microbiol. 2000, 3 (2), 107–111.

(31) Alves, L.; Paixão, S. M. Fructophilic Behaviour of Gordonia Alkanivorans Strain 1B during Dibenzothiophene Desulfurization Process. N. Biotechnol. 2014, 31 (1), 73–79. 10.1016/j.nbt.2013.08.007.

(32) Ram, S.; Mitra, M.; Shah, F.; Tirkey, S. R.; Mishra, S. Bacteria as an Alternate Biofactory for Carotenoid Production: A Review of Its Applications, Opportunities and Challenges. J. Funct. Foods 2020, 67 (November 2019), 103867. 10.1016/j.jff.2020.103867.

(33) Kim, J. H.; Kim, S. H.; Yoon, J. H.; Lee, P. C. Carotenoid Production from N-Alkanes with a Broad Range of Chain Lengths by the Novel Species Gordonia Ajoucoccus A2T. Appl. Microbiol. Biotechnol. 2014, 98 (8), 3759–3768. 10.1007/s00253-014-5516-y.

(34) Takaichi, S.; Maoka, T.; Akimoto, N.; Carmona, M. L.; Yamaoka, Y. Carotenoids in a Corynebacterineae, Gordonia Terrae AIST-1: Carotenoid Glucosyl Mycoloyl Esters. Biosci. Biotechnol. Biochem. 2008, 72 (10), 2615–2622. 10.1271/bbb.80299.

(35) Reginald, S. S.; Etzerodt, M.; Fapyane, D.; Chang, I. S. Functional Expression of a Mo-Cu-Dependent Carbon Monoxide Dehydrogenase (CODH) and Its Use as a Dissolved CO Bio-Microsensor. ACS Sensors 2021, 6 (7), 2772–2782. 10.1021/acssensors.1c01243.

(36) Gill, C. O. The Solubility of Carbon Dioxide in Meat. Meat Sci. 1988, 22 (1), 65–71. 10.1016/0309-1740(88)90027-7.

(37) Armor, J. N. Influence of PH and Ionic Strength upon Solubility of NO in Aqueous Solution. J. Chem. Eng. Data 1974, 19 (1), 82–84. 10.1021/je60060a013.

