## Supplementary material for "Unlocking the potential of *Gordonia rubripertincta* in syngas fermentation for carbon monoxide bioconversion into carotenoids": blastp

**SUPPLEMENTARY INFORMATION**

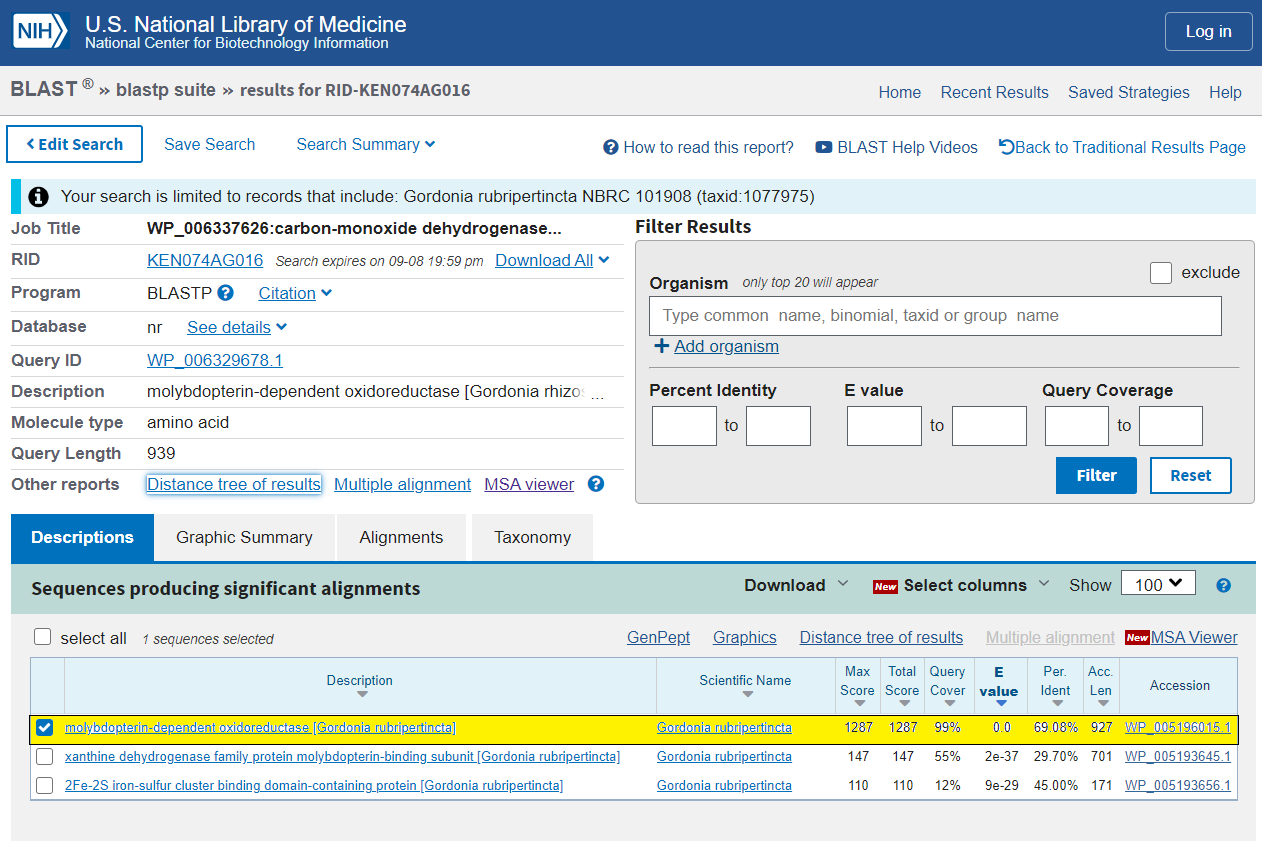

**Fig.S1. blastp result. The highlighted region shows the CODH in *G. rubripertincta* that is similar to CODH in *G. rhizosphera***

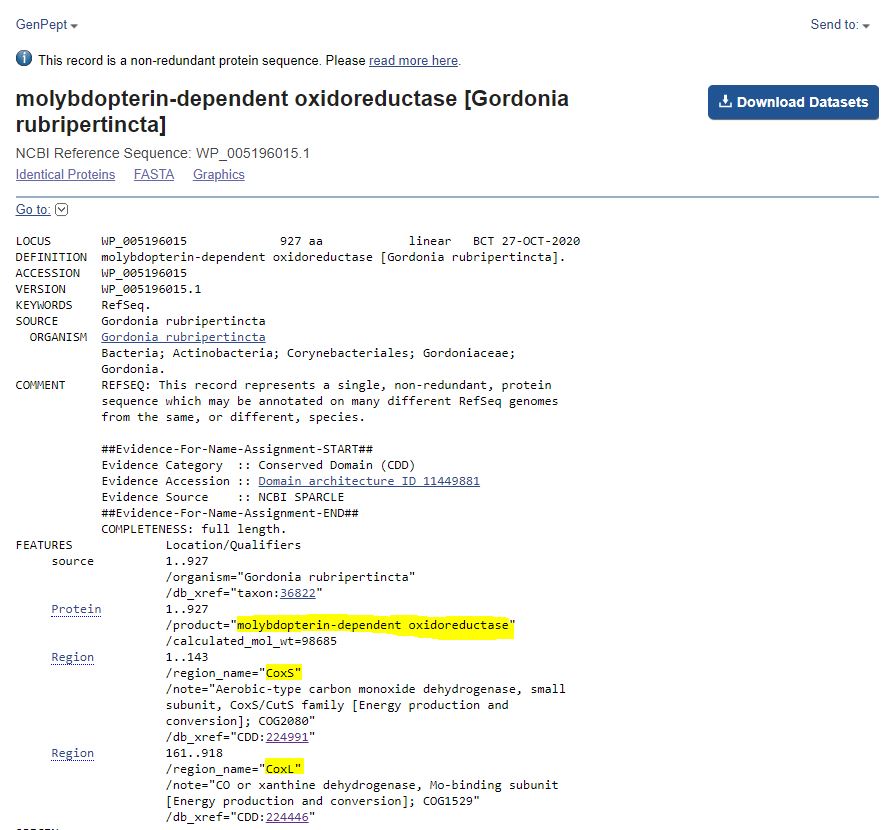

**Fig.S2. Details on the *G. rubripertincta* CODH enzyme**
